## Supplemental Figures for "Exploring the trade-off between deep-learning and explainable models for brain-machine interfaces"

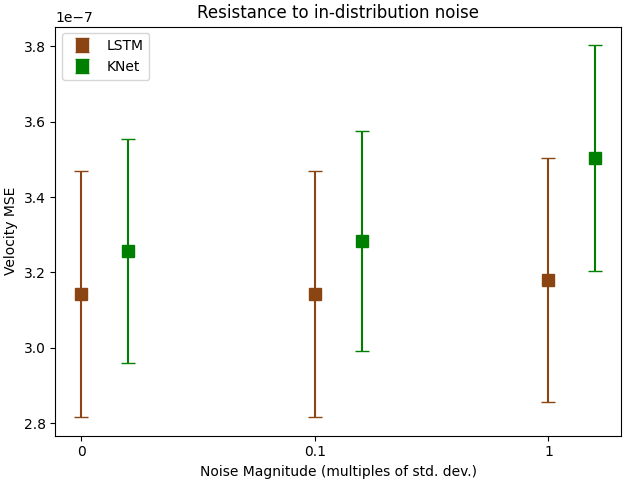

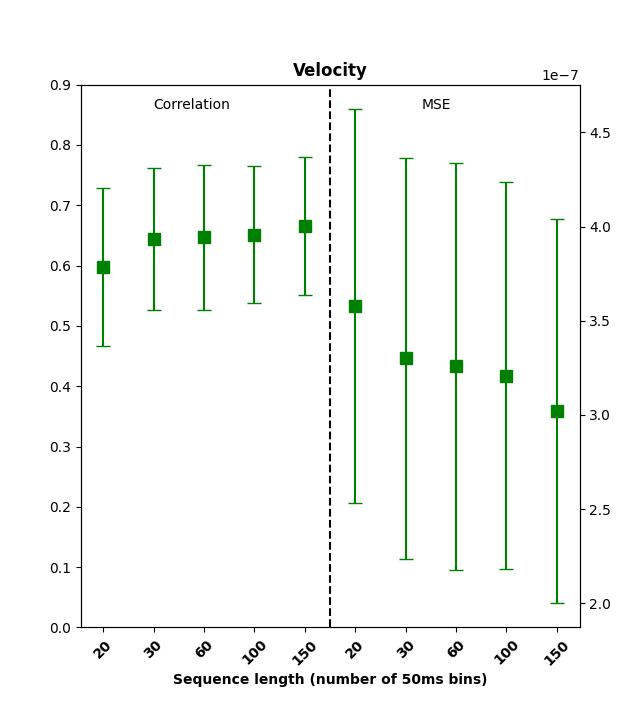

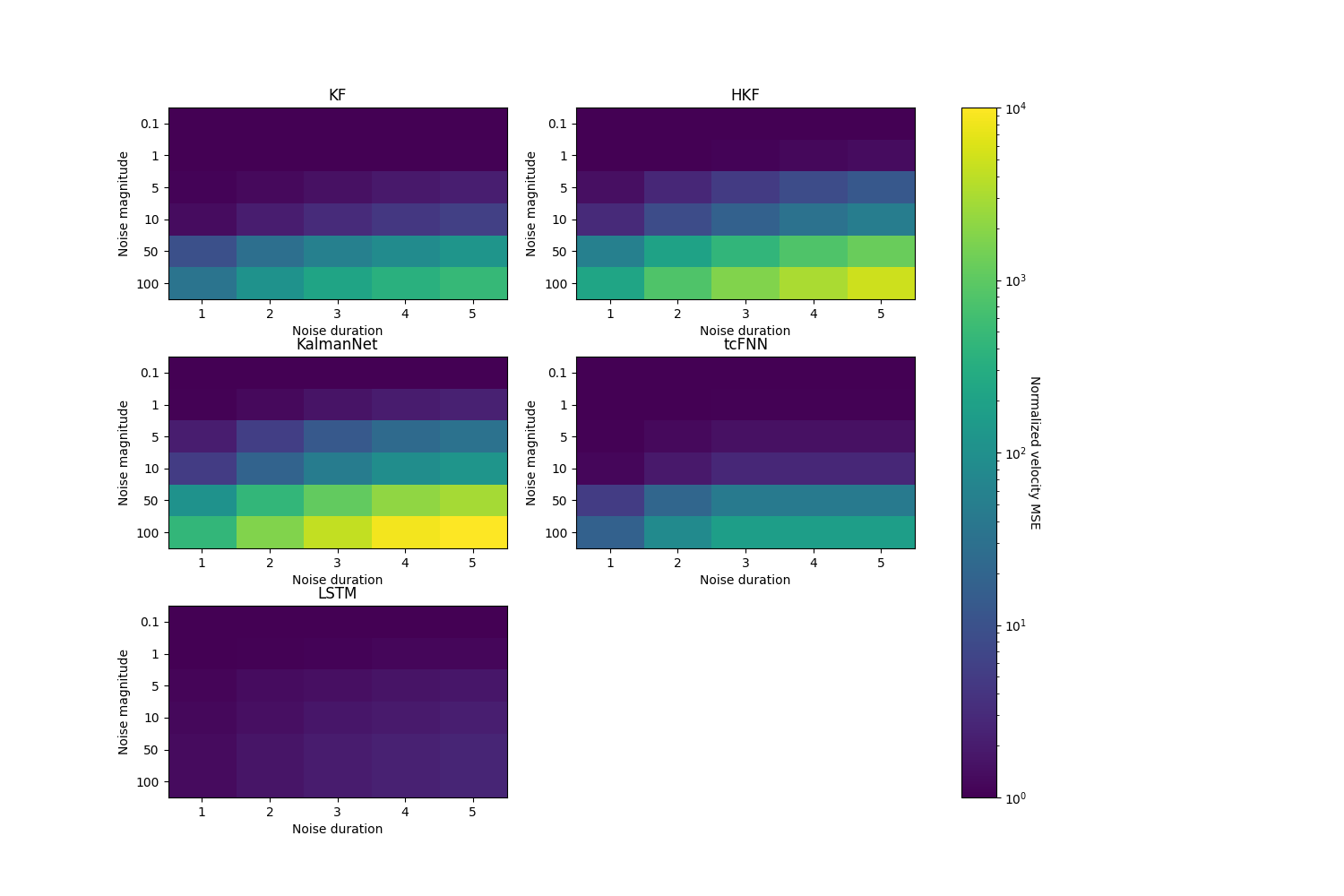

Supplemental Figure 1: **Resistance to in-distribution noise**. Offline velocity MSE for KalmanNet (green) and LSTM (brown) across n=13 days for both monkeys, with values of noise closer to those present in the training data. A noise of zero magnitude is equivalent to not adding noise (i.e., baseline shown on Figure 2).

Supplemental Figure 2: **Sensitivity analysis of sequence length during training**. Offline velocity correlation (left) and MSE (right) for KalmanNet, under different sequence lengths employed during training. The horizontal axis represents the number of 50ms bins; the one used throughout corresponds to 60, or equivalently, three seconds. Computed across all n=13 days for both monkeys.

Supplemental Figure 3: **Normalized MSE across models per noise magnitude and duration**. Full product of normalized velocity MSE across models. The logarithmic color bar on the right represents the MSE value for each combination of noise magnitude and duration, normalized to each model’s baseline performance (without noise).
